# Musicianship and melodic predictability enhance neural gain in auditory cortex during pitch deviance detection

**DOI:** 10.1101/2021.02.11.430838

**Authors:** D.R. Quiroga-Martinez, N. C. Hansen, A. Højlund, M. Pearce, E. Brattico, E. Holmes, K. Friston, P Vuust

## Abstract

When listening to music, pitch deviations are more salient and elicit stronger prediction error responses when the melodic context is predictable and when the listener is a musician. Yet, the neuronal dynamics and changes in synaptic efficacy underlying such effects remain unclear. Here, we employed dynamic causal modeling (DCM) to investigate whether the magnetic mismatch negativity response (MMNm)—and its modulation by context predictability and musical expertise—are associated with enhanced neural gain of auditory areas, as a plausible mechanism for encoding precision-weighted prediction errors. Using Bayesian model comparison, we asked whether models with intrinsic connections within primary auditory cortex (A1) and superior temporal gyrus (STG)—typically related to gain control—or extrinsic connections between A1 and STG—typically related to propagation of prediction and error signals—better explained magnetoencephalography (MEG) responses. We found that, compared to regular sounds, out-of-tune pitch deviations were associated with lower intrinsic (inhibitory) connectivity in A1 and STG, and lower backward (inhibitory) connectivity from STG to A1, consistent with disinhibition and enhanced neural gain in these auditory areas. More predictable melodies were associated with disinhibition in right A1, while musicianship was associated with disinhibition in left A1 and reduced connectivity from STG to left A1. These results indicate that musicianship and melodic predictability, as well as pitch deviations themselves, enhance neural gain in auditory cortex during deviance detection. Our findings are consistent with predictive processing theories suggesting that precise and informative error signals are selected by the brain for subsequent hierarchical processing.

**Significance statement:** In complex auditory contexts, being able to identify informative signals is of paramount importance. Such is the case of music listening, where surprising sounds play a fundamental role in its perceptual, aesthetical, and emotional experience. Crucially, surprising sounds in the pitch dimension are more easily detected and generate stronger cortical responses when melodies are predictable and when the listener is a musician. Using Dynamic Causal Modelling, here we show that such effects arise from a local increase in neural gain within auditory areas, rather than from passing of prediction and error signals between brain regions. Consistent with predictive processing theories, this suggests that the enhanced precision of auditory predictive models—through melodic predictability and musical training—up-regulates the processing of informative error signals in the brain.

## 1. Introduction

Surprising sounds in auditory sequences are perceived as more salient and generate stronger neural responses than expected sounds (Heilbron & Chait, 2018). The salience of surprising sounds and the responses they elicit depend on at least two factors: the musical expertise of the listener and the predictability of the auditory context. In music, for example, pitch deviants elicit a mismatch response—the mismatch negativity (MMN)—taken to reflect the update of the brain’s internal predictive model by prediction error (Vuust et al., 2012). Crucially, deviants are more easily detected and the amplitude of the MMN becomes larger, when melodies are predictable and when the listener is a musician (Quiroga-Martinez et al., 2019a). Yet, despite the growing empirical evidence (Garrido et al., 2013; Hsu et al., 2015; Kliuchko et al., 2019; Southwell & Chait, 2018; Tervaniemi et al., 2014), little is known about the fluctuations in effective connectivity and neuronal dynamics that mediate these phenomena.

The effect of context predictability on sound salience has been proposed to reflect the neural weighting of unexpected events by the precision afforded to sensory inputs (Quiroga-Martinez et al., 2019b; Ross & Hansen, 2016; Vuust et al., 2018). Such a precision-weighting mechanism would allow the brain to select informative sensory signals for further processing (Feldman & Friston, 2010; Friston et al., 2020; Hohwy, 2012). Research on attention has linked this selection to a modulation of postsynaptic gain in which the activity of neurons representing attended features and objects is enhanced (Garrido et al., 2018; Rabinowitz et al., 2015; Reynolds & Desimone, 1999). Such gain modulation likely arises from a change in the strength of intrinsic (i.e., within-region) connections controlling the excitability of brain areas (Auksztulewicz et al., 2017, 2018; Auksztulewicz & Friston, 2015), usually ascribed to NMDA receptor function and synchronous interactions between fast-spiking inhibitory interneurons and pyramidal cells. However, it remains unclear whether the same gain mechanisms operate when the salience of a sound is driven, not endogenously by selective attention, but rather exogenously by the predictability of successive stimuli.

The effect of musicianship on sound salience has also been suggested to rely on precision-driven mechanisms. Vuust et al. (2018) proposed that musicians possess a more precise predictive model of musical auditory signals than non-musicians, a view that has behavioral support (Hansen et al., 2016; Hansen & Pearce, 2014). If musicians have a fine-grained representation of musical tuning—which facilitates deviance detection and leads to enhanced MMN responses—the same precision-driven gain mechanisms above may also operate when sound salience is enhanced by musical expertise.

Here, we characterized the neuronal dynamics and effective connectivity underlying the salience of surprising musical sounds and its modulation by predictability and musical expertise. We employed dynamic causal modeling (DCM) of magnetoencephalography (MEG) data from a previous study investigating magnetic MMN responses (MMNm) in melodic sequences (Quiroga-Martinez et al., 2019a). In that experiment, musicians and non-musicians listened to highly predictable stimuli—a repeated four-note pattern—and less predictable stimuli—complex, less-repetitive melodies. We found that pitch deviants were more easily detected and generated larger MMNm responses in highly predictable compared to less predictable melodies, and in musicians compared to non-musicians.

We based the DCM analyses on an auditory network comprising bilateral primary auditory cortex (A1), bilateral superior temporal gyrus (STG), and the right frontal operculum (rFOP). We asked whether the MMNm, and its modulation by predictability and musical expertise, relied on changes in intrinsic connectivity within A1 and STG, as a plausible synaptic mechanism implementing precision-weighting of prediction error. We compared this to the alternative explanation that predictability and expertise modulate propagation of prediction and error signals through forward and backward extrinsic (i.e., between-regions) connectivity, which is typically associated with short-term plasticity, sensory learning and model updating. Thus, we contrasted models in which intrinsic, forward and/or backward connections were allowed to explain auditory evoked responses and their modulation as measured with MEG.

## 2. Methods

### 2.1. Participants

Twenty musicians and 20 non-musicians were included in the study. These participants are part of a larger group of 24 non-musicians and 26 musicians whose data have been analyzed and reported elsewhere (Quiroga-Martinez et al., 2019b, 2019a; Quiroga-Martinez, Hansen, et al., 2020). The four non-musicians and six musicians excluded were those for whom high-quality MRI images were not available, due to artifacts or abstaining from the MRI session. Musical expertise and musical competence (Table 1) were assessed with the Goldsmiths Musical Sophistication Index (GMSI) (Müllensiefen et al., 2014) and the Musical Ear Test (MET) (Wallentin et al., 2010). Musicians had significantly higher GMSI (*t*(31.1) = 14.82, *p* < .001) and were significantly better at differentiating melodies/rhythms than non-musicians, as indicated by the total scores on the MET (*t*(35.2) = 5.38, *p* < .001). See Quiroga-Martinez et al. (2020) for a more detailed report of musicianship measures. All participants gave informed consent and were paid 300 Danish kroner (approximately 40 euro) as compensation. The study was approved by the Regional Ethics Committee (De Videnskabsetiske Komitéer for Region Midtjylland in Denmark) and conducted in accordance with the Helsinki Declaration.

**Table 1.** Participants’ demographic and musical expertise information. Mean and standard deviation are reported.

|  | Musicians | Non-musicians |
| --- | --- | --- |
| Sample size | 20 | 20 |
| Female | 8 | 10 |
| Age | 24.2 ($\pm 3.09$) | 26.9 ($\pm 3.48$) |
| GMSI | 36.80 ($\pm 6.77$) | 10.65 ($\pm 4.05$) |
| MET | 84.05 ($\pm 6.98$) | 70.00 ($\pm 9.36$) |

### 2.2. Stimuli

In the experiment, we included conditions with high-predictability (HP) and low-predictability (LP) stimuli. HP stimuli comprised simple melodies consisting of a four-note repeated pitch pattern that has often been used in musical MMNm paradigms and is known as the Alberti bass (Vuust et al., 2011, 2012, 2016). LP stimuli consisted of a set of major and minor versions of six novel melodies, which had a much less repetitive internal structure and spanned a broader local pitch range than HP stimuli (Figure 1; see Supplementary file 1 in Quiroga-Martinez et al., 2019b for the full stimulus set). The predictability of these stimuli was measured in terms of Shannon entropy with IDyOM, a computational model of auditory expectations (Pearce, 2005). The corresponding analyses are reported in Quiroga-Martinez et al. (2019b) and revealed higher entropy values for the LP than the HP condition. Individual melodies were 32 notes long, lasted eight seconds, and were pseudo-randomly transposed between 0 and 5 semitones upwards. The presentation order of the melodies was pseudorandom within each condition. After transposition, the pitch-range of the LP condition spanned 31 semitones from B3 (F_0_ ≈ 247 Hz) to F6 (F_0_ ≈ 1397 Hz). LP melodies were transposed to two different octaves to cover approximately the same pitch range as HP melodies.

**Figure 1.**
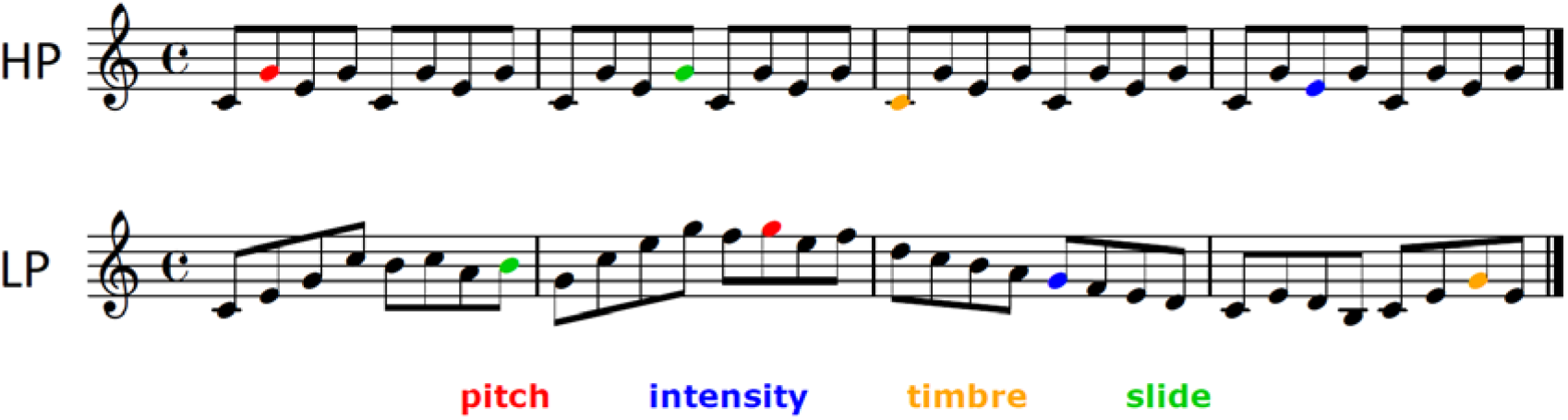
Example of the melodies used in the high predictability (HP) and low predictability (LP) conditions. Deviants are indicated with colors. Only pitch deviants were analyzed in this article.

For stimulus delivery, a pool of 31 standard piano tones was created with the “Warm-grand” sample in Cubase (Steinberg Media Technology, version 8). Each tone was 250 ms long, was peak-amplitude normalized and had 3-ms-long fade-in and fade-out to prevent clicking. No gaps between tones were introduced. Four types of deviants were pseudo-randomly introduced in the melodies, including pitch, timbre, intensity, and pitch-glide. Pitch deviants were created with Audition (Adobe Systems Incorporated, version 8) by raising the pitch of standard tones by 50 cents. Note that predictability in these stimuli was manipulated only in the pitch dimension by changing pitch-alphabet size and pitch repetitiveness, which resulted in pitch deviants exhibiting the strongest predictability effect in sensor-level analyses; see Quiroga-Martinez et al. (2019b, 2019a) for further details. Consequently, here we focused exclusively on pitch MMNm responses for the DCM analysis.

Each condition was presented in a separate group of three consecutive blocks, each around seven minutes long (i.e., HP-HP-HP/LP-LP-LP or LP-LP-LP/HP-HP-HP). Within each block, melodies were played one after the other without pauses. At the beginning of each block, a melody with no deviants was added to ensure a certain level of auditory regularity at the outset. One deviant per feature was introduced in each of the 144 melodies per block, amounting to a total of 144 deviants per feature in each condition. The position of each deviant was defined by segmenting the melody into groups of four notes (half a bar in Fig. 1), selecting some of these groups, and choosing randomly any of the four places within a group with equal probability. The order of appearance of the different types of deviants was pseudorandom, so that no deviant followed another deviant of the same feature. The selection of four-note groups was counterbalanced among melodies—under the constraints of a combined condition (i.e., melody and accompaniment) that was included to assess the predictive processing of simultaneous musical streams (see Quiroga-Martinez et al., 2019b, for further details). The analysis of the combined condition will be reported elsewhere. LP and HP conditions were counterbalanced across participants and always followed the combined condition.

### 2.3. Experimental Procedure

Participants received oral and written information, completed musical expertise questionnaires and put on MEG-compatible clothes. We then digitized their head shapes for co-registration with anatomical images and head-position tracking. During the recording, participants were sitting upright in the MEG scanner looking at a screen. Before presenting the musical stimuli, their individual hearing threshold was measured through a staircase procedure using a pure tone with a frequency of 1 kHz. The sound level was set at 60 dB above threshold. We instructed them to watch a silent movie of their choice, ignore the sounds and move as little as possible. Participants were informed there would be musical sequences playing in the background interrupted by short pauses so that they could take a break and adjust their posture. Sounds were presented through isolated MEG-compatible ear tubes (Etymotic ER•30). The recording lasted approximately 90 min, and the whole experimental session took between 2.5 and 3 hours, including consent, musical expertise tests, preparation, instructions, breaks, and debriefing.

### 2.4. MEG recording and preprocessing

Brain magnetic fields were recorded with an Elekta Neuromag MEG TRIUX system with 306 channels (204 planar gradiometers and 102 magnetometers) and a sampling rate of 1,000 Hz. Continuous head position information (cHPI) was obtained with four coils attached to the forehead and the mastoids. Offline, the temporal extension of the signal source separation (tSSS) technique (Taulu & Simola, 2006) was used to isolate signals coming from inside the skull employing Elekta’s MaxFilter software (Version 2.2.15). This procedure included movement compensation for all participants except two non-musicians, for whom continuous head position information was not reliable due to suboptimal placement of the coils. These participants, however, evinced reliable auditory event-related fields (ERFs), as verified by visual inspection of the amplitude and polarity of the P50(m) component. Electrocardiography, electrooculography, and independent component analysis were used to correct for eye-blink and heartbeat artefacts, employing a semi-automatic routine (FastICA algorithm and functions “find_bads_eog” and “find_bads_ecg” in MNE-Python) (Gramfort, 2013). Visual inspection of the rejected components served as a quality check.

Using the Fieldtrip toolbox (version r9093) (Oostenveld et al., 2011) in MATLAB (R2016a, The MathWorks Inc.), epochs comprising a time window of 400 ms after sound onset were extracted and baseline-corrected with a pre-stimulus baseline of 100 ms. Epochs were then low-pass-filtered with a cut-off frequency of 35 Hz and down-sampled to a resolution of 256 Hz. For each participant, ERFs were computed by averaging the responses for the deviants and averaging a matched selection of an equal number of standards. The evoked responses were then converted to SPM format for the source localization and DCM analyses. Sensor-level statistical analyses of these data have been reported in Quiroga-Martinez et al. (2019b, 2019a; 2020). Given that we used the same preprocessed data reported in our previous studies—where planar gradiometers were combined by taking the root mean square of channel pairs—in this paper, we restricted our analyses to magnetometer data, as in Quiroga-Martinez et al. (2020).

### 2.5. Source localization and network structure

To identify the auditory networks underlying the processing of pitch MMNm responses, we localized the neural generators of the standard and deviant evoked responses. For this, we used Multiple Sparse Priors (Friston et al., 2008) implemented in SPM12 (version 7478). T1- and T2-weighted magnetic resonance anatomical images (MRI) were obtained—in a separate session for each participant—with a magnetization-prepared two rapid gradient echo (MP2RAGE) sequence (Marques et al., 2010) in a Siemens Magnetom Skyra 3T scanner. The two images were then combined, segmented, projected into MNI coordinates, and automatically co-registered with the MEG sensor array using digitized head shapes and preauricular and nasion landmarks. We verified the quality of the co-registrations by visual inspection. Lead fields were computed using a single-shell boundary element model with 20,484 dipoles (fine grid). For each participant, we extracted a volume reflecting the mean inverse solution from 0 to 300 ms after sound onset. The resulting images were then submitted to a group-level repeated-measures ANOVA with deviance (standard vs. deviant) and predictability (HP vs. LP) as factors. No distinction between musicians and non-musicians was made in these analyses. The resulting statistical parametric maps were corrected for multiple comparisons with random field theory and revealed a clear effect of deviance on source strength. There were two clusters, one in each hemisphere, encompassing five peaks: left (lA1, x = −50, y = −16, z = −4) and right Heschl’s gyrus (rA1; x = 46, y = −16, z = 0), left (lSTG; x = −58, y = −6, z = 6) and right (rSTG; x = 56, y = 2, z = −1) anterior superior temporal gyrus, and right frontal operculum (rFOP; x = 50, y = 4, z = 12). The coordinates of these peaks were used as spatial priors for the five nodes or sources of our DCM network.

Note that this network is very similar to classical DCM studies of MMN responses (Dietz et al., 2014; Garrido et al., 2008; Garrido, Kilner, Kiebel, et al., 2009; Schmidt et al., 2013) with two important differences. First, we include anterior STG (instead of posterior STG or planum temporale), which has been related to the processing of pitch sequences (Gander et al., 2019; Patterson et al., 2002). Second, we include the FOP, which is posterior to the inferior frontal gyrus node used previously. To ensure the source estimates in the FOP node were not an artifact of source leakage, we evaluated the evidence for models with and without this node using Bayesian model comparison.

### 2.6. Dynamic causal modeling

The aim of DCM (Friston et al., 2003; Moran et al., 2013) is to assess the evidence for different hypotheses about how observed neuronal activity is generated. These hypotheses are specified through a generative model in which the activity of neuronal populations evolves dynamically according to the structure and (state or condition dependent) connectivity of the network. To estimate the model parameters characterizing directed (effective) synaptic efficacy the predicted neuronal responses are projected (through a lead field) to sensor space for comparison with the recorded data, in an iterative process that employs Bayesian inference (David et al., 2006). This furnishes a posterior distribution over all model parameters (i.e., effective connectivity and condition-specific changes) and the marginal likelihood of the data in the form of a variational free energy bound on model evidence.

The generative model used here is based on the canonical microcircuit (Bastos et al., 2012), which includes four populations of neurons: Spiny stellate cells, inhibitory interneurons, and superficial and deep pyramidal cells (Fig. 2). Within a brain region or node (denoted by *i*), intrinsic communication between these populations results in overall inhibition. Between region, extrinsic excitatory forward connections project from superficial pyramidal cells in lower areas to spiny cells and deep pyramidal cells in higher areas, whereas extrinsic inhibitory backward connections project from deep pyramidal cells in higher areas, to superficial pyramidal cells and inhibitory interneurons in lower areas. Note that some excitatory connections result in inhibition of their targets, mediated by intermediate inhibitory synapses not modeled explicitly.

**Figure 2.**
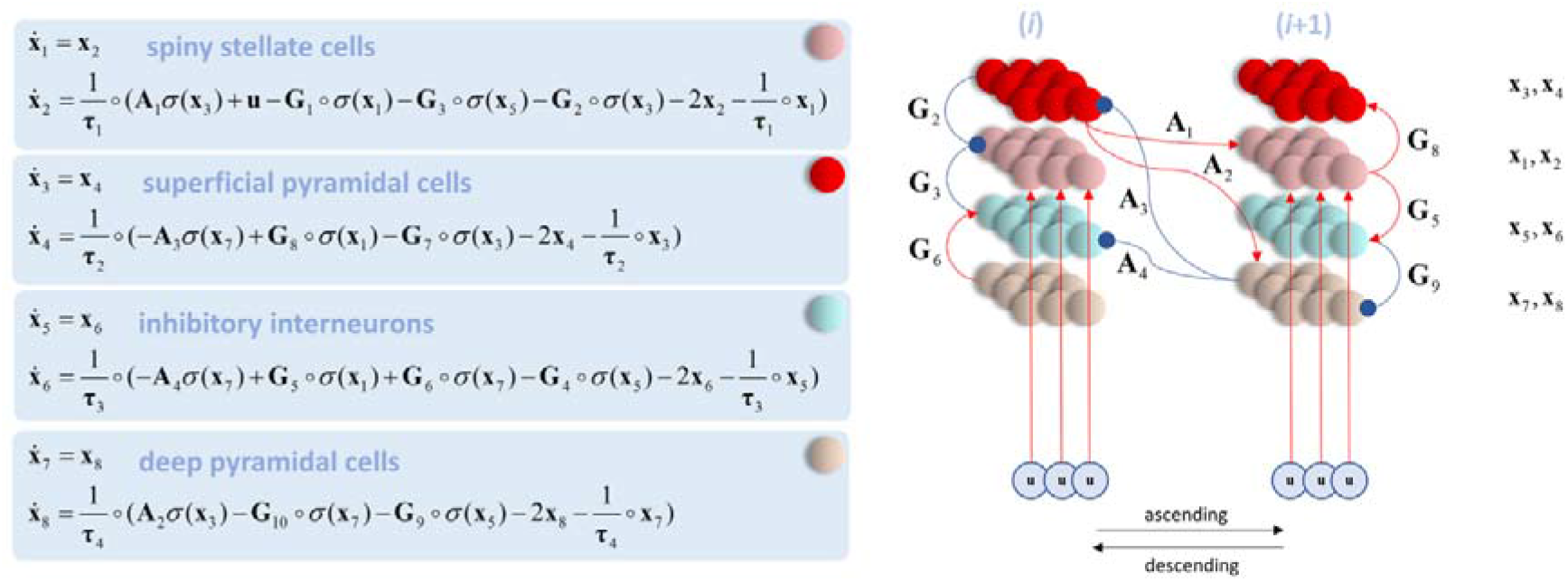
State equations (left) and generative model (right) based on the canonical microcircuit (Bastos et al., 2012). The activity of four populations of neurons (indicated by the **x** vectors and marked with different colors) evolves according to a set of coupled differential equations defined by the excitatory (red arrows) and inhibitory (blue arrows) connections in the network and a set of intrinsic (matrix G) and extrinsic (matrix A) synaptic weights. See main text for further details. Figure taken with permission from Parr et al. (2019).

The dynamics of the network are given by the set of coupled differential equations shown in Fig. 2. Here, the vector *x* represents the input current (even subscripts) and voltage (odd subscripts) of the corresponding populations of neurons. The matrices A and G respectively define the extrinsic and intrinsic connection strengths between neuronal populations. The time constants τ define the rate of change of the neuronal dynamics, whereas σ is a sigmoid activation function transforming voltages into firing rates. It is assumed that spiny cells receive input from lower regions denoted by **u.** The ∘ operator denotes element-wise product.

#### 2.6.1. Model structure and comparisons

We defined a network comprising five sources: rA1, lA1, rSTG, lSTG and rFOP. Forward connections were set from A1 to STG and from STG to FOP, whereas backward connections were set from FOP to STG and from STG to A1. The A1 nodes received simulated auditory thalamic input modelled with a temporal Gaussian bump function, peaking at 60 ms after sound onset. We used point dipoles to model each source. The prior location of each dipole was obtained from the source localization estimates above and the dipole orientation (and location) was optimized during model fitting. We assumed inter-hemispheric dipole symmetry, as warranted by the bilateral auditory generators.

In a first level (within subject) DCM analysis, we modeled the average evoked response from 0 to 300 ms after sound onset, separately for each of the two predictability conditions. We defined the standard sound as the baseline (reflected in the default connectivity matrix A) and allowed certain connections to change during processing of the deviant sound (as defined in a connectivity matrix B). Note that B-parameters effectively encode the difference between standards and deviants and, therefore, the modulation of effective connectivity that underwrites the MMN response. By switching on and off different B parameters, we were able to generate and test different hypotheses about how MMN responses are generated.

Thus, our model space included modulations of forward and backward connections between A1 and STG, modulations of intrinsic connections in A1 and STG, and different combinations of these. We also included models in which connections within, to, and from the FOP were switched on and off—to assess whether its contribution to source estimates was due to leakage from temporal areas, or whether it participated in the network dynamics. By switching on and off each of these families, we ended up with 2^4^ = 16 models: 15 grouped in the families intrinsic, forward, backward and FOP (Fig. 3b), and a null model in which no connections were modulated. Note that the families are not mutually exclusive such that a given model could belong to more than one family. For each participant and condition, we fitted the full model, which encompassed the modulation of all the B-parameters.

**Figure 3.**
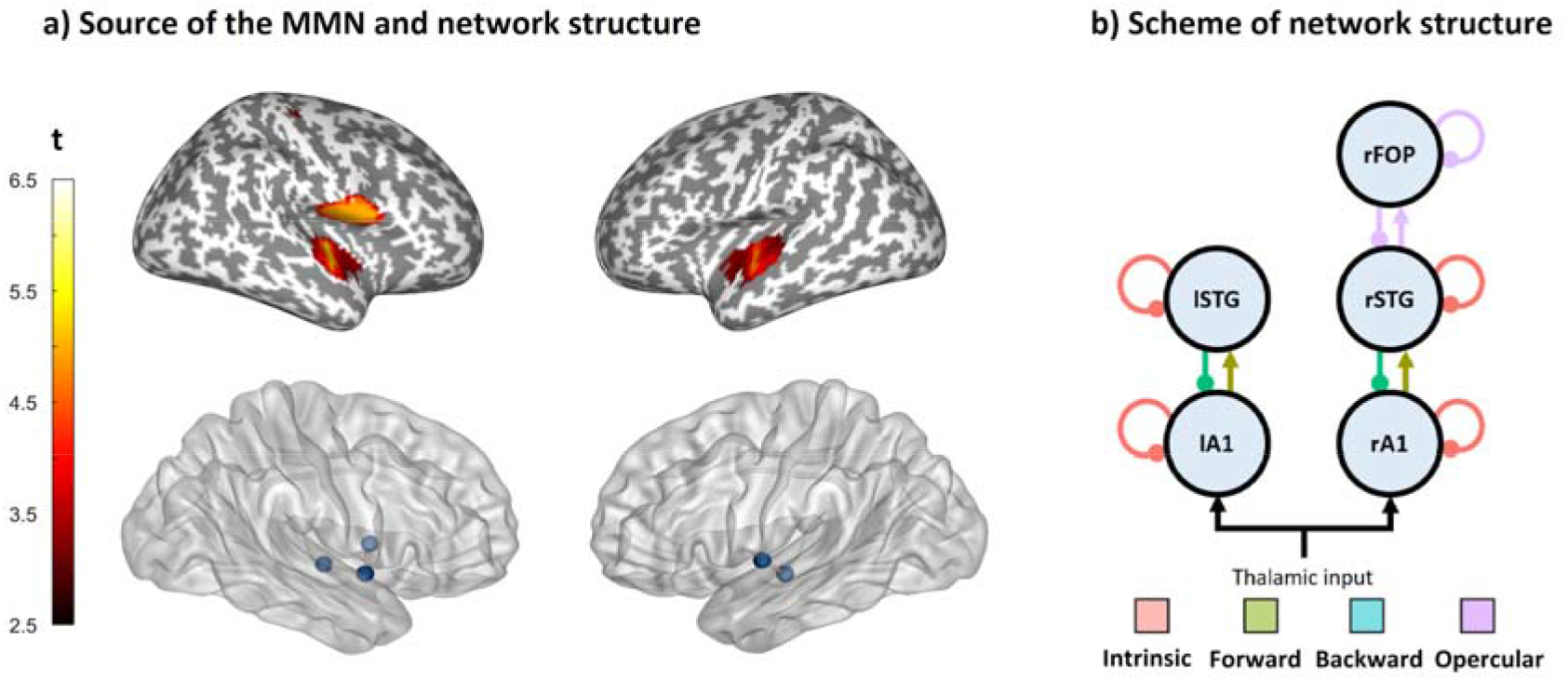
Structure of the network. a) Results of the source reconstruction (top) and prior location of the sources (bottom). b) Scheme of the network. The connections modulated in each of the four model families are indicated with colors. The combination of these families yielded a total of 2^4^ = 16 models, including a null model in which no connections were modulated. A1 = primary auditory cortex, STG = superior temporal gyrus, FOP = frontal operculum.

#### 2.6.2. Group-level analyses

First-level B-parameter estimates were submitted to a second level (between subject or group) analysis using Parametric Empirical Bayes (PEB), a technique in which the group-level variance is used as a hyperprior to constrain the random effects on first-level parameters (Zeidman et al., 2019). The second level model was a general linear model (GLM) that enabled us to assess the evidence that (a specific set of) B-parameters were modulated by stimulus deviance, melodic predictability, musicianship, or the predictability-by-musicianship interaction. The parameters of the second level model constitute an interaction between each factor of the GLM and each B-parameter of a DCM: for example, the effect of melodic predictability on the change in intrinsic connectivity within left A1.

All factors in the general linear model (i.e., design matrix) were mean-centered. To test our hypotheses, we assessed the evidence that the above factors had an effect on the B-parameters, by comparing the evidence for all models in which the factor is switched on, with that of all models in which it is switched off (including the null model). Note that an effect of stimulus deviance is simply a non-zero B-parameter, that is modelled with the constant term in the general linear model. The above analysis was repeated for a series of planned comparisons for each family of first level DCMs, i.e., prespecified hypotheses about the connections mediating the mismatch responses.

In a complementary analysis, we used Bayesian model averaging to estimate the effects parameterized by the second level model, i.e., the posterior density over model parameters weighted by the posterior probability of the models considered. Finally, in an exploratory analysis, we used Bayesian model reduction (Friston et al., 2016), which performs a “greedy” search over all possible models—including those outside our hypothesis space—by beginning with a full model that includes all the above effects, then pruning away redundant parameters (that are not necessary to account for the data and just add to model complexity). The (log) probability that a B-parameter is modulated by a factor is given as the difference in log evidence between second level models with and without the effect in question.

## 3. Results

### 3.1. Effective connectivity underlying the MMN

Compared to standard sounds, deviant sounds reduced the inhibitory intrinsic connections in A1 and STG and inhibitory backward connections from STG to A1. This is reflected in the high posterior probabilities (> .90) of “backward” and “intrinsic” families (Fig. 4a, first column). In contrast, the “forward” family has a much lower posterior probability (.32). The family-wise analysis also showed a low posterior probability for the “opercular” family, suggesting that modulations of frontal opercular activity are unlikely to explain the MMN responses observed in this experiment.

**Figure 4.**
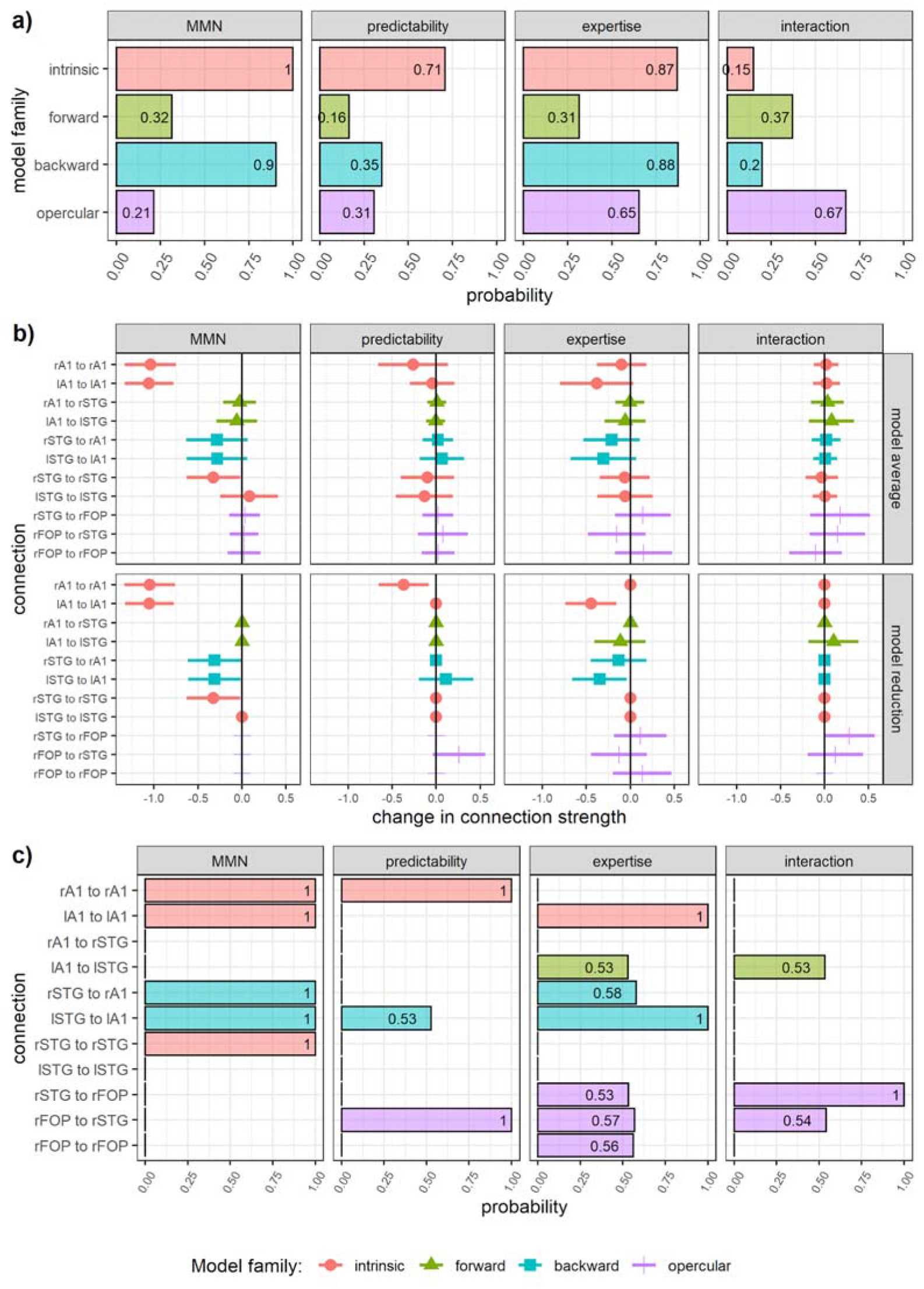
Results of group-level DCM analyses. (a) Posterior probability of the modulation of B-parameters in each family. Note that families are not mutually exclusive and therefore the sum of their probabilities may exceed 1. (b) Parameter estimates after Bayesian model reduction (top) and averaging (bottom), corresponding to the modulation of connection strengths by sound deviance (MMN), melodic predictability, musical expertise, and the predictability-by-expertise interaction. In model averaging, parameter values are weighted by the posterior probability of the models in the hypotheses space. In contrast, in exploratory model reduction, the evidence for parametric effects is obtained from an exhaustive search across all possible model configurations, in which effects with little evidence are pruned away (i.e., set to 0). Error bars represent 95% credible intervals. (c) Posterior probability of the modulation of each parameter after Bayesian model reduction.

The results of Bayesian model averaging, and Bayesian model reduction were largely consistent with the results of the planned Bayesian family comparisons. 95% credible intervals excluded zero in the case of intrinsic connections (rA1, lA1 and rSTG) for both model averaging and reduction, and in the case of backward connections (rSTG to rA1 and lSTG to lA1) for model reduction only (Fig. 4b, first column). Furthermore, intrinsic, and backward parameters had high posterior probabilities (> .99) after Bayesian model reduction (Fig. 4c, first column). The modulation of these connections by stimulus deviance implies a disinhibition and a corresponding increase in the excitability or gain, of A1 and rSTG contributions to the MMN.

To assess the accuracy of the DCM results, we compared the observed sensor-level data with those predicted by the model. Fig. 5 and 6 show that the predicted data match the observed data well, reproducing the topography of magnetic fields and the larger responses for predictable compared to less predictable melodies and for musicians compared to non-musicians, especially in the left hemisphere.

**Figure 5.**
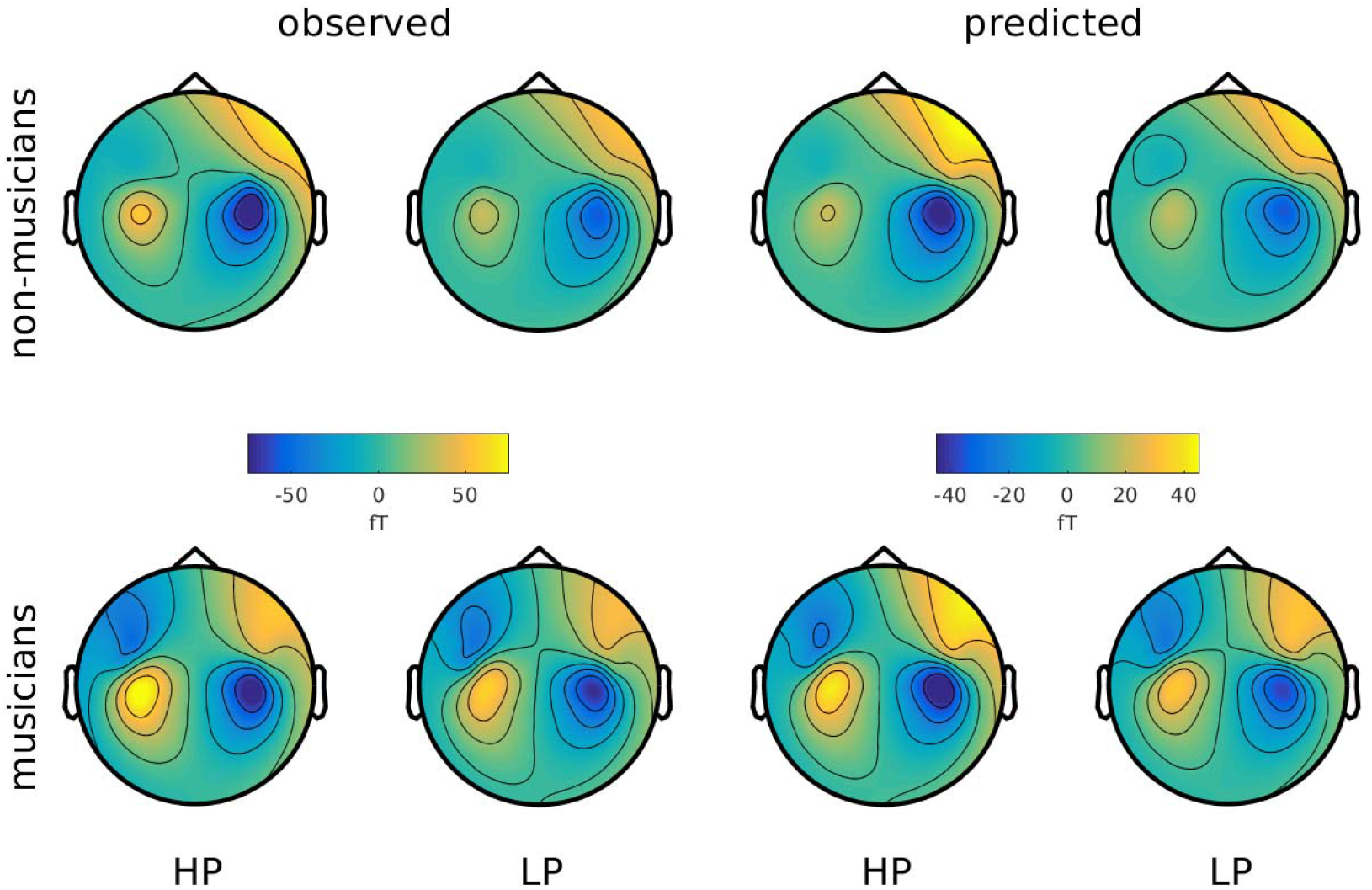
Topography of grand average MMNm responses (difference between deviants and standards) from 170 to 210 ms after sound onset, as observed in the experiment and predicted by DCM. HP = High predictability, LP = Low predictability.

**Figure 6.**
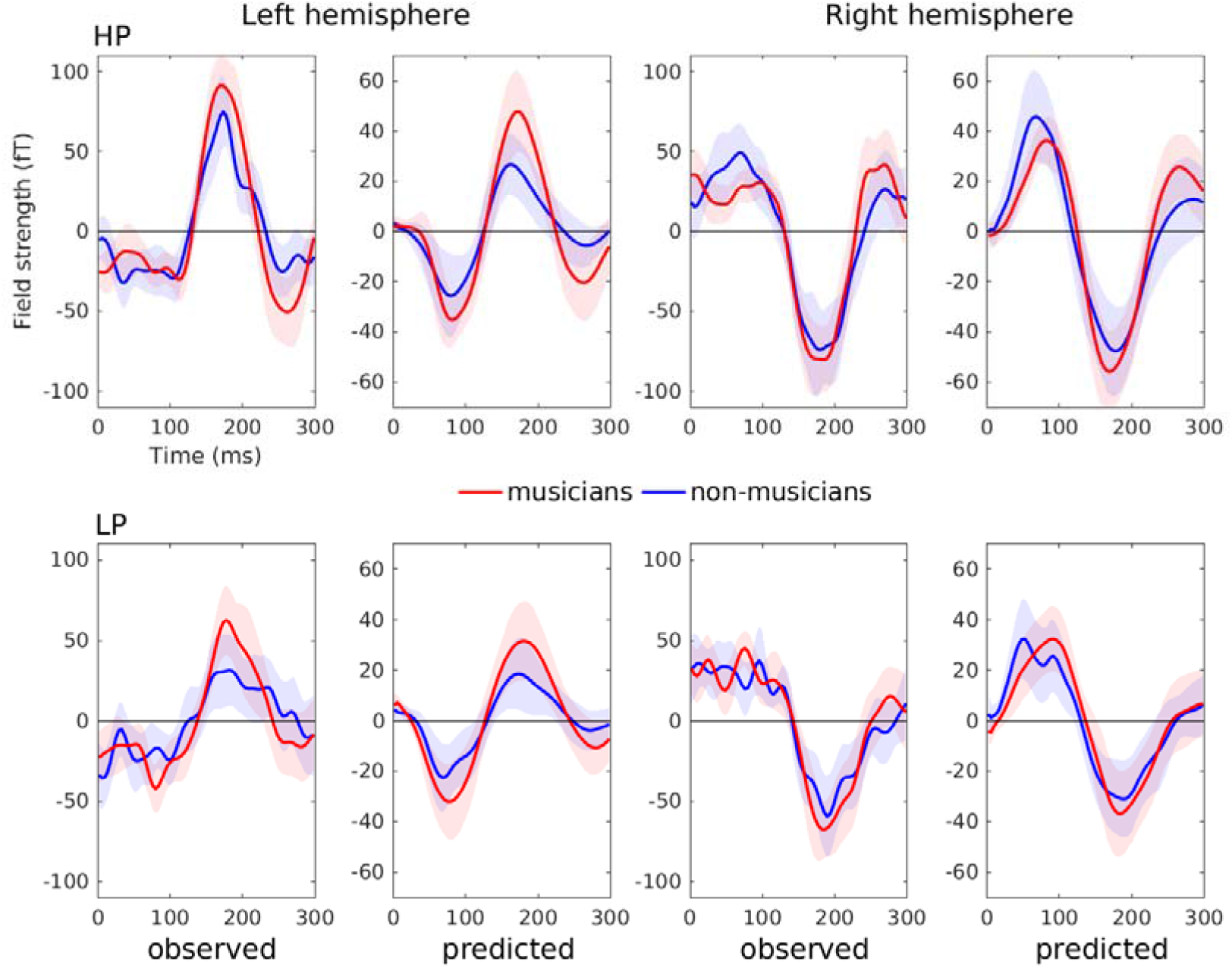
Event-related field of the MMNm (difference between deviants and standards) for each condition, group, and hemisphere, as observed in the experiment and predicted by DCM. The time courses correspond to the average of representative left (1611, 1621, 0231, 0241) and right (2411, 2421, 1331, 1341) auditory channels. Shaded areas depict 95% confidence intervals. HP = high predictability, LP = low predictability.

### 3.2. Effect of melodic predictability

We found reduced intrinsic connectivity in rA1 for HP compared to LP melodies, as shown by the relatively high probability of the “intrinsic” family (.71) (Fig. 4a, second column). In contrast, the posterior probabilities of forward (.16), backward (.25) and opercular (.31) families were much lower. The rA1 parameter had a high probability (> .99, Fig. 4c, second column) and its corresponding 95% credible interval excluded 0 (Fig. 4b, second column) after Bayesian model reduction. Furthermore, the exploratory model reduction also indicated the strengthening of the connection from rFOP to rSTG, as shown by the high probability of this effect (> .99, Fig. 4c, second column). However, this was not reflected in the probability of the opercular family (Fig. 4a, second column), so suggesting that this modulation was highly specific to the backward opercular connection within the right hemisphere and did not reflect a general contribution of the FOP.

### 3.3. Effect of musical expertise

We observed a decrease in the strength of intrinsic and backward inhibitory connections— i.e., disinhibition—in musicians compared to non-musicians, as shown in the high probability of “intrinsic” (.87) and “backward” (.88) families (Fig. 4a, third column). Posterior probabilities for forward (.31) and opercular (.65) families were low and moderate. 95% credible intervals excluded 0 in the case of the intrinsic lA1 connection for both model reduction and averaging, and in the case of the backward lSTG to lA1 connection for model reduction only (Fig. 4b, third column). Moreover, the probabilities of these two parameters were high after model reduction (> .99, Fig 4c, third column).

### 3.4. Interaction between predictability and expertise

We did not observe strong evidence for a predictability-by-expertise interaction, as reflected in the low posterior probabilities of intrinsic (.15), forward (.37) and backward (.20) families (Fig. 4a, fourth column). However, the probability of the opercular family was moderate (.67) and exploratory model reduction suggested a strengthening in the connection from rSTG to rFOP for musicians in the predictable condition. For this parameter, the credible interval excluded 0 (Fig 4b, fourth column, bottom panel) and the posterior probability was high (>.99, Fig. 4c, fourth column).

## 4. Discussion

In this study, we found that the MMN responses elicited by surprising sounds in music listening—and the effects of predictability and expertise on these responses—rest on disinhibition of auditory areas, as indicated by reduced intrinsic connectivity within A1 and STG and backward connectivity from STG to A1. This supports the notion that neural gain, as a plausible mechanism for mediating precision-weighted prediction error, underlies the salience of surprising sounds and the strength of the neural responses they generate.

### 4.1. Connectivity patterns underlying the MMN

The main contributors to the MMN were modulations of intrinsic connectivity within bilateral A1 and rSTG, and bilateral backward connectivity from STG to A1. Reduced intrinsic connectivity implies an increase in the excitability of neural populations and has been interpreted as a salience-related enhancement of neural gain in response to deviant sounds (Auksztulewicz et al., 2017, 2018; Auksztulewicz & Friston, 2015). In other words, mistuned sounds may attract attentional resources and thus be prioritized at the earliest stages of auditory cortical processing.

That we found a reduction in top-down inhibition from secondary to primary auditory areas and a lack of modulation of forward connections contrasts with most previous studies in which both forward and backward connections show oddball-related effects (Auksztulewicz & Friston, 2015; Chennu et al., 2016; Garrido et al., 2007, 2008; Garrido, Kilner, Kiebel, et al., 2009; Lumaca et al., 2020; Schmidt et al., 2013). From a predictive coding perspective (Friston, 2005; Garrido, Kilner, Stephan, et al., 2009; Huang & Rao, 2011), forward and backward communication between brain areas reflects the update of predictive models by prediction error. Thus, the reason why backward connections were weakened, and forward connections remained unchanged might be that there was little model updating (i.e., sensory learning) elicited by the deviants. This, in turn, might indicate that out-of-tune sounds are heard as occasional, attention-grabbing “wrong” notes, rather than structurally novel events demanding a change in the current predictive model. In other words, a tone in a melody could still be predicted and recognized, even when it is saliently out-of-tune. A similar view has been proposed by Koelsch et al. (2018), who suggest that, in typical musical MMN designs, the higher-order predictive model is so strong that deviant sounds elicit prediction error responses that do not get resolved at higher processing levels. As the authors put it, deviant sounds “fall on deaf ears” and “keep knocking on the door” (p. 6).

Another finding that differs from previous research is the lack of involvement of frontal areas in the generation of the MMN, as indicated by the low probability of the opercular family. This is consistent with the lack of frontal generators previously reported for the same dataset (Quiroga-Martinez et al., 2019a) and in a recent fMRI study using simple musical stimuli (Lumaca et al., 2020). This suggests that the opercular peak found in the present source-level statistical analyses may reflect source leakage. Thus, the acoustic deviations introduced (i.e., out-of-tune sounds) may have been resolved at low-level processing stages in the temporal lobe, without engaging frontal areas typically involved in the sequential processing of sounds (Koelsch et al., 2002, 2009). This may have been reinforced by the fact that participants were instructed to ignore the sounds and watch a film instead.

Interestingly, an EEG study using the same stimuli as here found that MMN responses were similarly modulated by melodic predictability in participants with congenital amusia (a condition that disrupts pitch processing) and controls (Quiroga-Martinez, Tillmann, et al., 2020). Since amusia most likely arises from reduced connectivity between temporal and frontal areas (Albouy et al., 2013; Peretz, 2016), this further indicates that the processes underlying the mistuning MMN and its modulation by predictability and musicianship are restricted to local auditory areas in the temporal lobe. Note, however, that exploratory Bayesian model reduction indicated that connections between rFOP and rSTG may have been modulated by predictability and the predictability-by-expertise interaction. This could indicate that, as sounds become more salient, higher-order brain areas are engaged. However, further research is needed to properly assess this claim and dissociate it from source leakage.

### 4.2. Enhancement of neural gain in predictable melodies

The strength of intrinsic (inhibitory) connections in rA1 was reduced in predictable compared to less predictable melodies. Such connectivity changes may thus underlie the stronger MMN response observed for the former. This is consistent with the hypothesis that the stimulus-driven increase in predictive precision enhances the sensitivity to upcoming sensory signals, thus providing evidence for the role of gain modulation in precision weighting of prediction error. This effect was found only in the right hemisphere, which may reflect the fact that musical pitch processing in the general population is predominantly right-lateralized (Albouy et al., 2020; Zatorre et al., 2002).

### 4.3. Left-lateralized gain enhancement in musicians

Compared to non-musicians, musicians showed disinhibition in lA1 and backward connections from lSTG to lA1. This indicates that the stronger MMN response in this group might rest on the same gain enhancement mechanism found for the effect of predictability. In turn, this suggests that both phenomena could be framed as enhancements in predictive precision. In previous work, we have proposed the terms “stimulus-driven” and “expertise-driven” to refer to these two types of uncertainty reduction (Quiroga-Martinez et al., 2019a). Thus, here we show that, although they seem to affect prediction error responses independently, stimulus-driven, and expertise-driven uncertainty reduction might rely on similar underlying changes in effective connectivity. Furthermore, these results agree with the enhanced responses previously found in musicians for violations of pitch-related regularities such as interval, contour, musical tuning, and pitch patterns (Boh et al., 2011; Fujioka et al., 2004; Herholz et al., 2009; Koelsch et al., 1999; Tervaniemi et al., 2014; Vuust et al., 2012).

Interestingly, the expertise-related gain enhancement was left lateralized, which adds to a collection of left-hemisphere specific effects linked to musical expertise (Ellis et al., 2013; Elmer et al., 2013; Limb et al., 2006; Ono et al., 2011; Tervaniemi et al., 2011; Vuust et al., 2005). Considering the hemispheric specialization for temporal vs. spectral information and for music vs. language (Albouy et al., 2020; Zatorre et al., 2002), this could mean that musicians’ pitch processing involves a finer temporal evaluation of the sounds and more explicit lexical knowledge of musical structure.

### 4.4. A plausible mechanism for precision-weighted prediction error

Taken together, our results suggest that the modulation of gain in auditory areas may underlie the weighting of prediction error responses by uncertainty (Clark, 2013; Feldman & Friston, 2010; Hohwy, 2012), where uncertainty corresponds to unpredictable melodic contexts and a lack of musical expertise in the current experimental design. Effects on intrinsic connectivity are not necessarily explained by short or long-term synaptic plasticity, but rather on modulations of synaptic efficacy through acetylcholine or other classical neuromodulatory neurotransmitters (Auksztulewicz & Friston, 2016; Baldeweg et al., 2006). Changes in synaptic efficacy may also be mediated by fast synchronous interactions, involving spiking inhibitory interneurons equipped with NMDA receptors (Schmidt et al., 2013). Consistent with our results, the precision weighting of prediction errors has been cast as reflecting unexpected uncertainty—i.e., a momentary change in the estimated predictability of the context induced by unexpected events—which has been associated with modulations of pupil diameter and the neuromodulator norepinephrine (Bianco et al., 2020; Dayan & Yu, 2006; Zhao et al., 2019). Thus, a plausible hypothesis is that the enhanced excitability of auditory cortex in response to deviant sounds has its origins in neuromodulation — and concomitant changes in synchronous gain. Note, however, that we also found evidence for a reduction of backward (inhibitory) connectivity, suggesting that the observed effects were, at least in part, mediated by changes in the sensitivity to top-down afferents from cortical sources higher in the auditory hierarchy. These changes, nonetheless, are quantitatively smaller than those in intrinsic connections and further contribute to the disinhibition of A1, thereby facilitating gain enhancement. Future research should aim to disentangle the specific contribution of neuromodulation and changes in synaptic efficacy to gain control in A1.

## 5. Conclusion

In this study, we characterized the neuronal dynamics and changes in synaptic efficacy underlying the salience of pitch deviants and its modulation by melody predictability and musical expertise during music listening. Using DCM, we found that musicianship and predictability, as well as deviance itself, increased neural gain in primary auditory cortex through a reduction in the strength of intrinsic (inhibitory) connectivity in A1 and STG. The MMN effect was also associated with reduced backward connectivity from STG to A1. Gain modulation in primary auditory cortex was right-lateralized in the case of predictability and left-lateralized in the case of musical expertise. Our findings are consistent with predictive processing theories suggesting that precise and informative error signals are prioritized by the brain for subsequent hierarchical processing. Furthermore, they suggest that the ability to contextualize sensory processing in musicianship and predictable sensory streams relies on similar neuronal gain mechanisms.

## Acknowledgements and funding

We wish to thank the project initiation group, namely Christopher Bailey, Torben Lund and Dora Grauballe, for their help with setting up the experiments. We also thank Nader Sedghi, Massimo Lumaca, Giulia Donati, Ulrika Varankaite, Giulio Carraturo, Riccardo Proietti, and Claudia Iorio for assistance during MEG recordings. Finally, we thank Hella Kastbjerg for checking the English language of this manuscript. The Center for Music in the Brain is funded by the Danish National Research Foundation (DNRF 117). EH is funded by Action on Hearing Loss (PA_25). NCH received funding from the European Union’s Horizon 2020 research and innovation programme under the Marie Skłodowska-Curie grant agreement No 754513 and The Aarhus University Research Foundation. KJF was funded by a Wellcome Trust Principal Research Fellowship (Ref: 088130/Z/09/Z). The funders had no influence on the scientific content of this article.

## Open practices

Code and data for the study are available at https://doi.org/10.17605/osf.io/bdr73

## CRediT authorship contribution statement

David R. Quiroga-Martinez: Conceptualization, Methodology, Software, Formal analysis, Data curation, Writing - original draft, Visualization, Investigation. Niels Chr. Hansen: Conceptualization, Methodology, Writing - original draft. Andreas Højlund: Conceptualization, Methodology, Software, Writing - original draft, Supervision. Marcus T. Pearce: Software, Formal analysis, Writing - original draft. Emma Holmes: Supervision, Writing - original draft. Elvira Brattico: Conceptualization, Supervision, Writing - original draft. Karl Friston: Supervision, Writing - original draft. Peter Vuust: Conceptualization, Methodology, Supervision, Writing - original draft, Funding acquisition.

